# Interplay between brassinosteroids and TORC signaling in Arabidopsis revealed by integrated multi-dimensional analysis

**DOI:** 10.1101/2021.02.12.431003

**Authors:** Christian Montes, Ching-Yi Liao, Trevor M Nolan, Gaoyuan Song, Natalie M Clark, Hongqing Guo, Diane C Bassham, Yanhai Yin, Justin W Walley

## Abstract

Brassinosteroids (BR) and Target of Rapamycin Complex (TORC) are two major processes coordinating plant growth and stress responses. BRs function through a signaling pathway to extensively regulate gene expression and TORC is known to regulate translation and autophagy. Recent studies revealed that these two pathways crosstalk, but a system-wide view of their interplay is still missing. Thus, we performed transcriptome, proteome, and phosphoproteome profiling of Arabidopsis mutants with altered levels of either BIN2 or RAPTOR1B, two key players in BR and TORC signaling, respectively. We found that perturbation of BIN2 or RAPTOR1B levels affects a common set of gene-products involved in growth and stress responses. Additionally, we performed Multiplexed Assay for Kinase Specificity (MAKS), which provided a system-wide view of direct BIN2 substrates. Furthermore, phosphoproteomic data was used to reconstruct a kinase-signaling network and to identify novel proteins dependent on BR and/or TORC signaling pathways. Loss of function mutants of many of these proteins led to an altered BR response and/or modulated autophagy activity. Altogether, these results provide genome-wide evidence for crosstalk between BR and TORC signaling and established a kinase signaling network that defines the molecular mechanisms of BR and TORC interactions in the regulation of plant growth/stress balance.

## Main

When plants respond to stress, specific molecular and cellular processes are triggered, and growth is often compromised. Therefore, well-coordinated crosstalk of different signaling pathways is fundamental for a successful response (Huot et al., 2014; Verma et al., 2016). The growth-promoting hormone brassinosteroid (BR) is critical for this balance. Plants with perturbed BR biosynthesis or signaling exhibit altered growth and response to stresses (Clouse et al., 1996; Li et al., 1996; Guo et al., 2013; Nolan et al., 2017a, 2017b; Gruszka, 2018; Ye et al., 2019; Nolan et al., 2020). The Glycogen Synthase Kinase 3 (GSK3)-like kinase BIN2 is a pivotal negative regulator of BR signaling (Li and Nam, 2002; Kim et al., 2009). In the absence of BRs, BIN2 phosphorylates the BES1/BZR1 family of transcription factors (TFs), reducing their protein levels, DNA binding, and promoting cytoplasmic sequestration by 14-3-3 proteins, thereby preventing the activation of downstream BR response genes (Ryu et al., 2007; Gampala et al., 2007; Ryu et al., 2010; Yin et al., 2002). Besides regulating BES1 and BZR1, increasing evidence positions BIN2 as a hub for stress and growth balance regulation (Cai et al., 2014; Cho et al., 2014; Hu and Yu, 2014; Youn and Kim, 2015; Jiang et al., 2019; Ye et al., 2019; Li et al., 2020a; Nolan et al., 2020; Li et al., 2020b).

In Arabidopsis, the target of rapamycin regulatory complex (TORC) is a vital regulator integrating nutrient and energy sensing into cell proliferation and growth (Xiong and Sheen, 2014; Fu et al., 2020). The complex is comprised of TOR kinase, LST8, and RAPTOR. RAPTOR interacts with and recruits substrates to the complex for phosphorylation by TOR (Hara et al., 2002; Yang et al., 2013). Two *RAPTOR* homologs, *RAPTOR1A* and *RAPTOR1B*, have been found in Arabidopsis with *RAPTOR1B* being the predominantly expressed copy (Anderson et al., 2005; Deprost et al., 2005). Activation of TORC signaling induces the expression of ribosomal proteins, increases protein translation, stimulates photosynthesis, and upregulates (transcriptionally and translationally) plant growth-promoting genes (Ren et al., 2012; Xiong et al., 2013; Dong et al., 2015; Van Leene et al., 2019; Scarpin et al., 2020). Conversely, TORC actively represses autophagy, a central recycling system of cytoplasmic components and whose regulation is essential for rerouting nutrients and other raw materials when needed for plant growth, development, or stress responses (Noda and Ohsumi, 1998; Pu et al., 2017; Marshall and Vierstra, 2018).

When plants encounter stress, autophagy is often triggered, and growth-promoting pathways such as BR or TORC signaling need to be dampened (Nolan et al., 2017a; Liao and Bassham, 2020). To enable this balanced regulation of plant growth and stress responses, hormonal pathways such as auxin (Li et al., 2017; Schepetilnikov et al., 2017) and BRs (Zhang et al., 2016; Vleesschauwer et al., 2018) can influence or be affected by TORC activity. Increasing evidence points towards TORC-regulated autophagy as a crucial interaction point between BRs and TORC signaling when controlling this balance. For example, activation of TORC signaling promotes BR response by stabilizing BZR1, likely preventing its autophagy-driven degradation (Zhang et al., 2016). Additionally, BIN2 knock-down lines exhibit reduced sensitivity to TOR inhibitors AZD8055 (AZD) and KU63794 (Xiong et al., 2017). Furthermore, S6K2 can phosphorylate BIN2 in a TOR-dependent manner. However, the mechanism and biological implications of this interaction are not clear. Moreover, BIN2 has been shown to phosphorylate ubiquitin receptor DSK2 to facilitate its interaction with ATG8 and promote BES1 degradation via selective autophagy (Nolan et al., 2017b).

BIN2 and TORC regulate plant response to environmental changes via phosphorylation, exerting molecular changes at many different levels (i.e., changes in gene transcription or protein activity) (Guo et al., 2013; Bozhkov, 2018; Van Leene et al., 2019; Liao and Bassham, 2020; Nolan et al., 2020). Therefore, understanding the molecular connection between BR and TORC signaling across different levels of gene expression is necessary to unravel the interplay between these pathways. Furthermore, despite BIN2 being intensively studied, proteome-wide identification of BIN2 substrates is lacking. Here, we present a comprehensive multi-omic profiling detailing transcriptome, proteome, and phosphoproteome changes that occur in mutants with altered levels of BIN2 or the TORC subunit RAPTOR1B. We complement these global in vivo profiles with proteome-wide identification of direct BIN2 substrates using a Multiplexed Assay for Kinase Specificity (MAKS). A significant overlap was found in the transcripts, proteins, and phosphosites whose accumulation is dependent on BIN2 and RAPTOR1B. We reconstructed a kinase-signaling network and used it to identify novel genes whose mutant lines showed either altered growth in response to BR and/or levels of autophagy. Together, these studies further our understanding of the dynamic interplay between BR and TORC signaling.

## Results

### Comprehensive multi-omics profiling of *bin2* and *raptor1b* mutants

We designed a multi-omics experiment to identify novel components dependent on BR and/or TORC signaling. We performed transcriptome, proteome, and phosphoproteome profiling on rosette leaves of 20-day old wild-type (WT), *bin2D* (gain-of-function), *bin2T (bin2 bil1 bil2* triple loss of function mutant), and *raptor1b* plants. We quantified transcript levels using 3’ QuantSeq and measured protein abundance and phosphorylation state using two-dimensional liquid chromatography-tandem mass spectrometry (2D-LC-MS/MS) on Tandem Mass Tag (TMT) labeled peptides (McAlister et al., 2012; Hogrebe et al., 2018; Song et al., 2018a) (Fig. 1a). From these samples, we detected 23,975 transcripts, 11,183 proteins, and up to 27,887 phosphosites from 5,675 phosphoproteins (Fig. 1b and Supplementary Data Set 1).

**Fig. 1.**
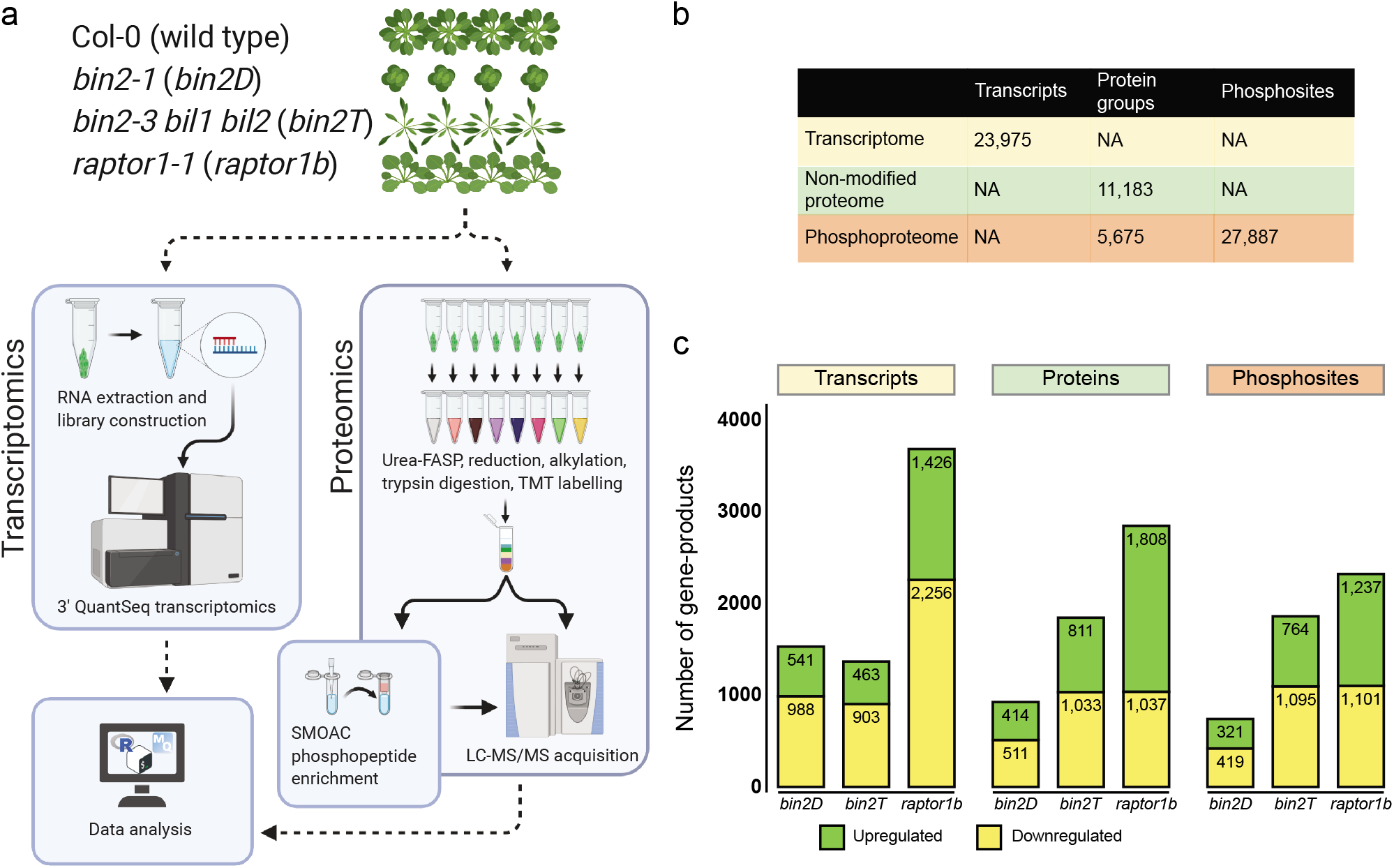
Experimental design, workflow, and data overview. **a**, Schematic representation of the multi-omics processing pipeline for *bin2* and *raptor1b* mutants. **b**, number of total detected transcripts, proteins, phosphoproteins, and phosphosites. **c**, differentially expressed transcripts, proteins, and phosphorylated amino acids for each analyzed mutant compared to WT.

We found 5,653 transcripts and 4,001 proteins that were differentially expressed (DE) in at least one mutant when compared against WT (Fig. 1c and Supplementary Fig. 1a and 2a). Gene ontology (GO) analysis of DE transcripts and proteins in *bin2D, bin2T*, and *raptor1b* mutants showed enrichment of many terms from similar processes including growth, hormones, stimuli sensing, and stress (Supplementary Figs. 1 and 2). These results suggest that the biological processes being affected at both protein and transcript level remain closely linked. Furthermore, the enriched terms among *bin2* and *raptor1b* DE transcripts and proteins are consistent with their known roles in growth/stress balance and hormonal crosstalk (Supplementary Data Set 2 and 3).

Our phosphoproteomics analysis identified 4,153 differentially phosphorylated sites in at least one mutant (Fig. 1c and Supplementary Fig. 3a). GO analysis for potential BIN2 target proteins (i.e., those with increased phosphorylation in *bin2D* or decrease phosphorylation in *bin2T*) revealed enrichment of terms related to plant growth and development, as well as response to stress and defense, processes in accordance with known BR functions. In addition, response to BR, abscisic acid, and auxin terms were also significant, highlighting once more the close relationship between BIN2 activity and these hormones. Transcriptional regulation-related terms were significantly enriched in the *bin2D* dataset, consistent with BIN2’s well documented regulatory activity upon TFs (Supplementary Fig. 3b,c and Supplementary Data Set 4). Finally, we assessed GO enrichment for proteins with decreased phosphorylation in *raptor1b*. We found that most of the enriched terms were related to growth, autophagy, starvation, auxin, and BR response. This is consistent with the known biological role of RAPTOR and suggests a cross-regulation between BR and TORC pathways via phospho-signaling (Supplementary Fig. 3d and Supplementary Data Set 4).

### BIN2 and RAPTOR1B regulate a core set of common gene-products

Next, we compared the DE transcripts we identified with previously reported transcriptomic data for BR and TORC. We found a significant overlap between the DE transcripts in *bin2* mutants and transcripts that respond to brassinolide (BL) treatment (Wang et al., 2014); 40.2% of *bin2D (p* = 6.14e-45) and 36.8% *bin2T (p* = 2.85e-24) responded to BL treatment. Interestingly, 30.9% of *raptor1b* DE transcripts (p = 6.55e-22) overlapped with BL responsive transcripts (Supplementary Fig. 4a). We also compared our data to the transcriptome profile of 10-day-old Arabidopsis seedlings treated with mock or AZD8055 (AZD), a specific TOR kinase inhibitor (Dong et al., 2015). We observed a significant overlap (*p* = 1.03e-15) between the mis-expressed transcripts in *raptor1b* and transcripts DE in AZD treated plants. Similar to the overlap between *raptor1b* and BL treatment, AZD responsive transcripts exhibited a significant overlap with transcripts DE in both *bin2* mutants. Specifically, 25.5% of transcripts DE in *bin2D (p* = 5.85e-38) and 26.2% of transcripts DE in *bin2T (p* = 1.18e-35) were also AZD responsive (Supplementary Figure 4b).

The reciprocal enrichment of BL-responsive transcripts in *raptor1b* and AZD (i.e., TORC) responsive transcripts in *bin2* mutants prompted us to investigate whether there are common gene-products being regulated by both BIN2 and RAPTOR1B. We found significant overlaps in DE geneproducts between *raptor1b* and *bin2D* as well as between *raptor1b* and *bin2T* mutants, at all three levels (transcriptome, proteome, and phosphoproteome). For our transcriptomic data we found that 25.3% of genes DE in the *bin2D* mutant (p = 2.08e-15) were also DE in the *raptor1b* mutant and 30.0% of genes DE in *bin2T (p* = 2.01e-30) were also DE in *raptor1b*. Interestingly, from our proteomic analyses we found that 55.8% of proteins DE in *bin2D (p* = 1.02e-24) and 50.6% of those DE in *bin2T (p* = 3.97e-26) were also DE in the *raptor1b* mutant. Finally, we found that 26.7% of *bin2D* DE phosphosites (p = 2.04e-16) and 21.9% of *bin2T* DE phosphosites (p = 8.64e-16) were DE in *raptor1b* (Fig. 2a). Since we found a significant overlap between our *raptor1b* mutant and both *bin2* mutants, we next investigated the biological processes being jointly affected. GO analysis on the proteins DE in both *raptor1b* and *bin2D* or *bin2T* mutants showed significant enrichment of stress terms (response to toxic substance, response to heat, response to cold, response to oxidative stress) and growth-related processes (photosynthesis, glucose metabolic process, ribosome assembly and translation) (Fig. 2b and Supplementary Data Set 5). These results reveal that BIN2 and RAPTOR1B affect a common core set of gene-products. Additionally, GO analysis suggest an involvement of these genes in processes related to growth and stress responses.

**Fig. 2.**
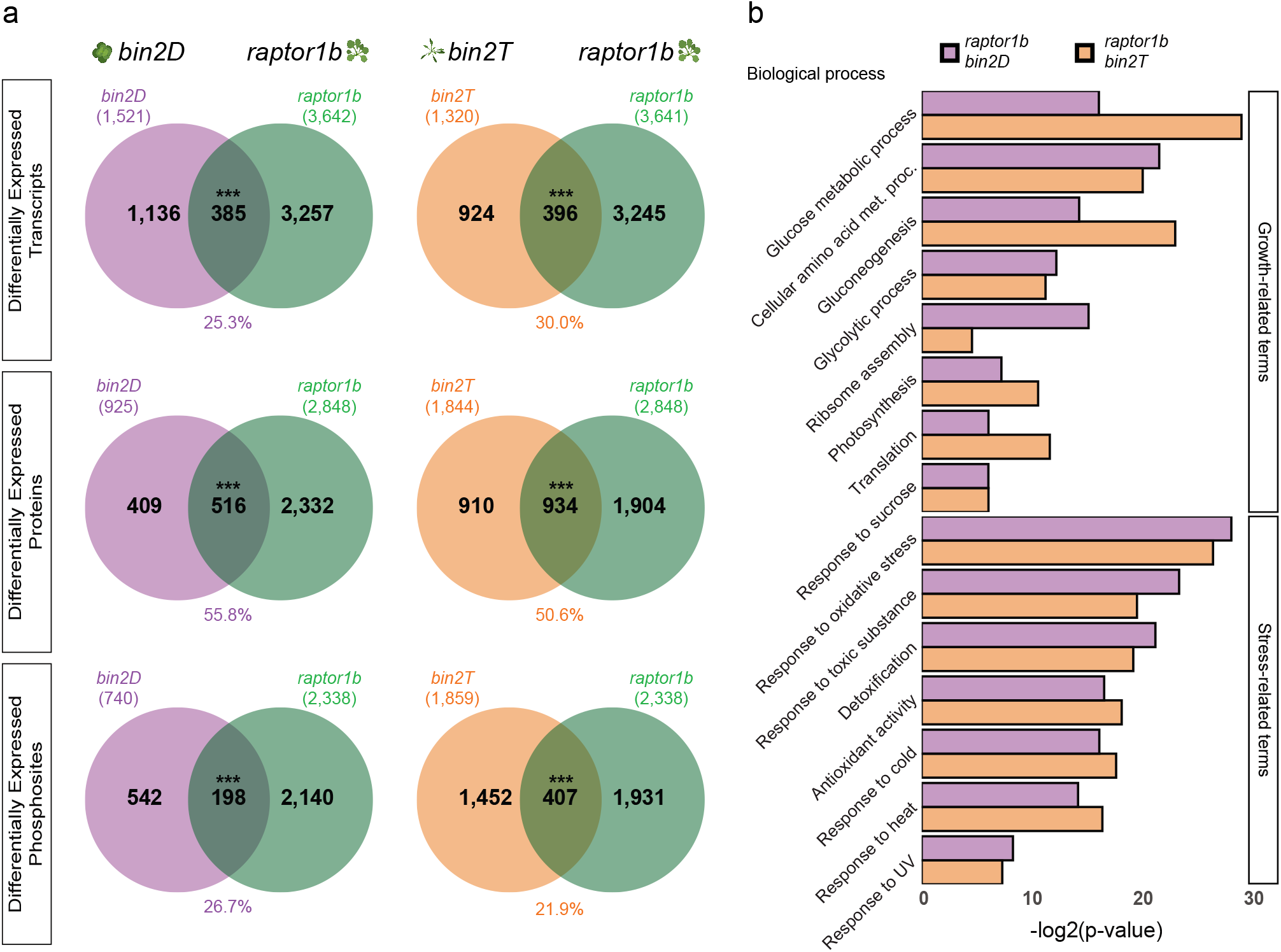
BIN2 and RAPTOR1b affect a common core set of common gene-products. **a**, Venn diagram showing overlap of differentially expressed transcripts (upper), proteins (middle), or phosphosites (lower) between *raptor1b* and *bin2D* (left) or bin2T (right) mutants. Statistical significance was assayed using hypergeometric test, assessing only gene-products present in both datasets (*** *p* < 0.001). **b**, Significantly enriched GO biological processes among proteins whose expression is being affected concomitantly by *raptor1b* and either *bin2D* (purple) or bin2T (orange). GO significance calculated using Fisher’s exact test, significance cutoff was established as q-value < 0.1.

### Phosphoproteomic analysis of *bin2* mutants shows enrichment of BIN2 direct targets

Because BIN2 is a kinase, we hypothesized that phosphosites increased in the *bin2D* gain-of-function mutant or decreased in *bin2T* may be direct BIN2 substrates. To test this hypothesis, we generated a proteome-wide dataset of BIN2 direct targets using the multiplexed assay for kinase specificity (MAKS) (Brumbaugh et al., 2014; Jayaraman et al., 2017) (See Methods for details). We quantified a total of 10,375 phosphosites accounting for 3,628 phosphoproteins from this assay (Supplementary Data Set 1b). The obtained phosphoproteome was heavily skewed toward increased phosphosites (Fig. 3a), with 1,343 phosphosites increased following incubation with GST-BIN2 (Supplementary Data Set 1b). Among proteins with increased phosphorylation we observed YDA and BSK1, two known BIN2 targets (Kim et al., 2012; Sreeramulu et al., 2013). To evaluate this set of phosphorylation sites as BIN2 kinase-substrates, we performed motif enrichment analysis and found a significant enrichment of the well-known GSK3 motif “S/T-x-x-x-S/T” (Fiol et al., 1987; Youn and Kim, 2015) among the increased phosphosites (*p* = 2.01e-25, Fig. 3b and Supplementary Data Set 6a). Additionally, another highly enriched motif found in the analysis was “S/T-P”, which is reported as a motif for GSK3, CDK, and MAPK families (Amanchy et al., 2007; Lin et al., 2015) (Supplementary Data Set 6a). Some previously unreported length variations of the GSK3 motif were also significantly enriched (i.e., S/T-x-x-S/T, S/T-x-S/T, and S/T-S/T, Supplementary Data Set 6a). These results support the robustness of our BIN2 kinase dataset.

**Fig. 3.**
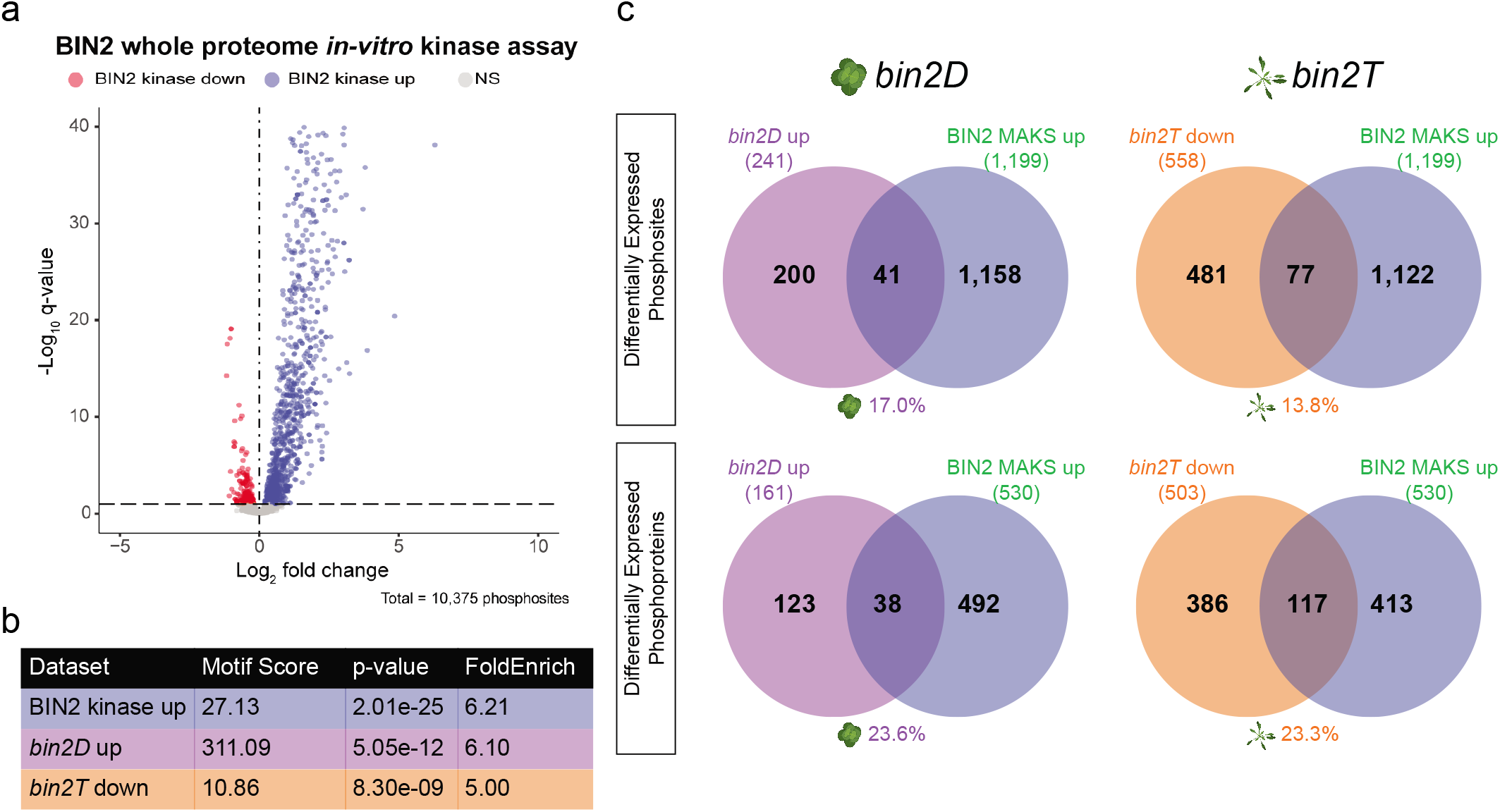
Phosphoproteomic analysis on BIN2 mutants shows significant enrichment of BIN2 direct targets. **a**, Volcano plot of phosphorylation sites from a Multiplexed Assay for Kinase Specificity (MAKS) on Arabidopsis leaf protein extracts incubated with recombinant GST or GST-BIN2. Significantly increased phosphosites are colored blue whereas decreased phosphosites are red (q < 0.1). **b**, De novo motif enrichment analysis showed high enrichment for the GSK3 motif on BIN2-related phosphoproteomic datasets. Motif score and FoldEnrich values are calculated by motifeR, while *p*-value was calculated using hypergeometric testing. **c**, Venn diagrams show overlap between BIN2 direct targets (i.e., those upregulated in BIN2 MAKS) and phosphosites (upper) or phosphoproteins (lower) upregulated in *bin2D* (left) or downregulated in *bin2T* (right) mutants. Numbers below each Venn diagram represent the overlapping percent for that mutant (purple = *bin2D*, orange = *bin2T*). Phosphosite overlaps were calculated using a 20 amino acid window, centered on the differentially regulated phosphosite (for details see methods section). Statistical significance was calculated using hypergeometric testing (* *p* <0.05, ** *p* < 0.01).

We next assessed the prevalence of BIN2 direct targets present in our profiling of *bin2D* and *bin2T* mutants. For this, we first performed motif enrichment analysis on phosphosites either upregulated in *bin2D* or downregulated in *bin2T*. A significant enrichment for the GSK3 motif was found in both *bin2D* (*p* = 5.05e-12) and *bin2T* (*p* = 8.30e-09) phosphosites (Fig. 3b and Supplementary Data Set 6b,c). Next, we looked at the overlap with our BIN2 direct targets identified in the MAKS experiment. For *bin2D*, 17.0% (41/241; *p* = 6.53e-03) of the total differentially up-regulated phosphosites and 23.6% (38/161; *p* = 2.58e-03) of the upregulated phosphoproteins were also BIN2 direct targets. For *bin2T*, 13.8% (77/558; *p* = 5.04e-02) of the downregulated phosphosites and 23.3% (117/503; *p* = 5.80e-06) of the downregulated phosphoproteins were also part of our BIN2 direct substrate list (Fig. 3c and Supplementary Data Set 7). These results indicated that a subset of the BIN2 dependent phosphosites identified by in vivo mutant profiling may be direct BIN2 substrates.

### Kinase-signaling network inference on *bin2* and *raptor1b* mutants

Since both BIN2 and RAPTOR1B (TORC) participate in phosphorylation-based signaling, differentially accumulating phosphosites identified in our mutant profiling maybe direct or indirect targets of BIN2 and/or RAPTOR1B. With this in mind, we reconstructed the molecular relationships of these signaling networks. To do so, we used our data to infer a kinase-signaling network for each mutant (i.e., *bin2D, bin2T*, and *raptor1b*). To build these networks, we first inferred the activation state of kinases in our dataset. The activation loop (A-loop) is a well-conserved region inside the kinase domain whose phosphorylation is necessary for kinase activation (Adams, 2003; Ahiri, 2019). Thus, phosphosite intensity level of the A-loop can be used as a proxy for kinase activity quantification (Walley et al., 2013; Beekhof et al., 2019; Schmidlin et al., 2019). First, we performed a whole-proteome Arabidopsis in-silico A-loop prediction and were able to identify the region on 1,360 proteins (Supplementary Data Set 8a). Subsequently, we identified kinases whose A-loop phosphosite intensity was differentially regulated in at least one of the profiled mutants (Supplementary Fig. 5a). We found 27, 21, and 24 kinases exhibiting an altered activation state in the *bin2D, bin2T*, and *raptor1b* mutant, respectively (Supplementary Data Set 8b). Using this information, we inferred a kinase-signaling network by correlating phosphosite level with kinase activation state (Supplementary Fig. 5b). A network containing 4,138 nodes, representing 33 activated kinases and 2,284 target sites arising from 1,853 possible substrate proteins was obtained (Fig. 4a and Supplementary Data Set 9). To evaluate this kinase-signaling network, predicted BIN2 targets were obtained (i.e., nodes connected by edges directed outward of BIN2), and motif enrichment analysis was performed. As expected, the GSK3 motif was enriched among BIN2 targets (p = 1.96e-06). Additionally, the MAPK consensus motif P-X-[pS/pT]-P was overrepresented among MPK6 targets (*p* = 3.56e-16). Moreover, several variants of the proline-directed phosphorylation motif [pS/pT]-P, were significantly enriched among targets of MPK4 (*p* < 0.001), MPK10 (*p* < 0.001), and BIN2 (*p* < 0.001) (Amanchy et al., 2007; Lin et al., 2015; Rayapuram et al., 2021) (Fig. 4b and Supplementary Data Set 10). Furthermore, 66.7% (6/9) of known BIN2 targets reported in the literature and present in our network were correctly predicted as BIN2 targets. These results support the target prediction value of our inferred kinase signaling network.

**Fig. 4.**
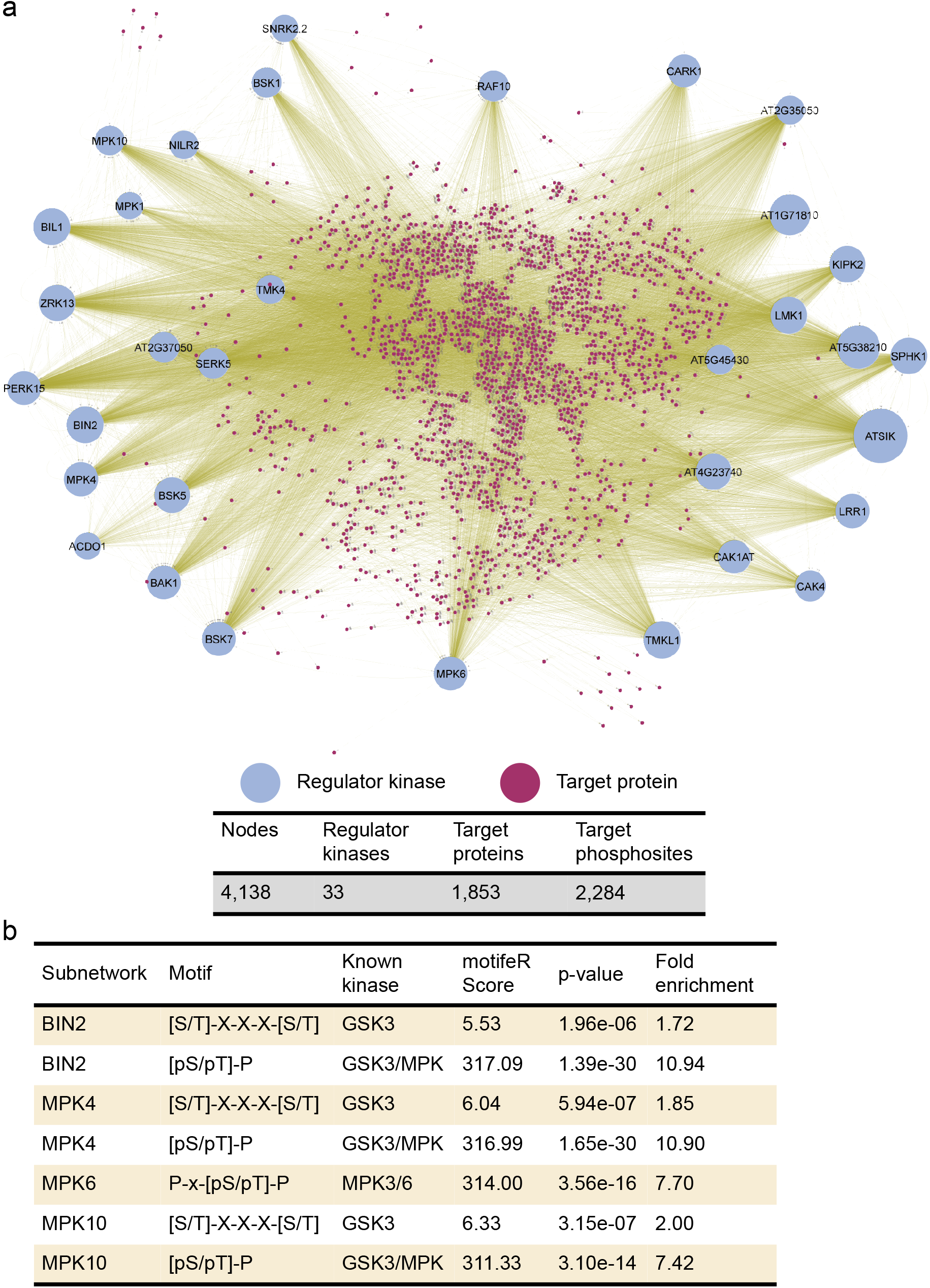
Kinase signaling network. **a**, A signaling network was inferred using phosphoproteomic data from *bin2D, bin2T*, and *raptor1b* mutants. Activated kinases are shown as named circles with their size representing the number of predicted targets (i.e., node outdegree). Target proteins are represented as small, purple circles. **b**, De novo motif enrichment analysis among predicted direct targets (i.e., node first neighbors) for BIN2, MPK4, and MPK10 showed high enrichment for the GSK3 motif ([S/T]-X-X-X-[S/T]) and GSK3/MPK3 motif ([pS/pT]-P). Analysis on MPK6 predicted direct targets showed significant enrichment of MPK3/6 motif (P-X-[pS/pT]-P). Enrichment analysis was done on a 14 amino acids window, centered on target phoshposites.

### Phosphoproteomics and kinase-signaling network reconstruction reveal proteins required for normal BR response and autophagy

After analyzing the phosphoproteome of our *bin2* and *raptor1b* mutants and predicting their signaling network, the next logical step was to determine whether these predictions identified proteins bridging both BR response and TORC-related autophagy. To assess this question, we enlisted those proteins being differentially phosphorylated simultaneously in either of the BIN2 mutants (i.e., *bin2D* or *bin2T*) and in *raptor1b* (Supplementary Fig. 6a,b). When selecting candidates, we focused our attention first on those proteins up-phosphorylated in *bin2D* and those down-phosphorylated in *bin2T* since this phosphorylation “directionality” could pinpoint those proteins direct or indirectly affected by BIN2 activity. We then fine-tuned this selection by keeping only those proteins exhibiting differential phosphorylation in *raptor1b* (Supplementary Fig. 6a,b). From this universe, we selected 42 candidate genes present in our kinase-signaling network (either as a target, an activated kinase, or both) and obtained mutants to assess their BR and autophagy phenotypes (Supplementary Data Set 11). Mutant lines for these candidate genes were tested for hypocotyl elongation in response to BL as a means to assess their sensitivity to BR. From the tested candidate genes, 33.4% (14/42) showed a significantly altered BL response (Fig. 5a and Supplementary Data Set 11a). To assess autophagy levels as a readout of TORC activity, a total of 28 candidate genes that showed significantly altered hypocotyl elongation in response to BL, exhibited decreased phosphorylation in *raptor1b* mutant, or both, were stained using the fluorescent dye monodansylcadaverine (MDC) (Floyd et al., 2015). The number of acidic vesicles as a proxy for the number of autophagosomes occurring in the root elongation zone was then assessed by microscopy (Floyd et al., 2015). Twenty genes (71.4% of assayed candidate genes) showed consistently increased autophagy levels when mutated (Fig. 5a and Supplementary Data Set 11b). We further confirmed the autophagy phenotype on selected mutants by examining GFP-ATG8e marker, which labels autophagosomal membranes, by transient expression in protoplasts obtained from mutant lines or in stable GFP-ATG8e transgenic lines in the mutant backgrounds (Xiong et al., 2007; Floyd et al., 2015). From this confirmation process, 100% of assayed mutants showed increased autophagy when compared to WT (Supplementary Data Set 11c). Additionally, mutants for the six genes without significant increase in autophagy under normal conditions were subjected to sucrose starvation and stained with MDC. All six mutants showed a reduced induction of autophagy upon starvation when compared to WT, with four showing a complete loss of induction (Fig. 5a and Supplementary Data Set 11d).

**Fig. 5.**
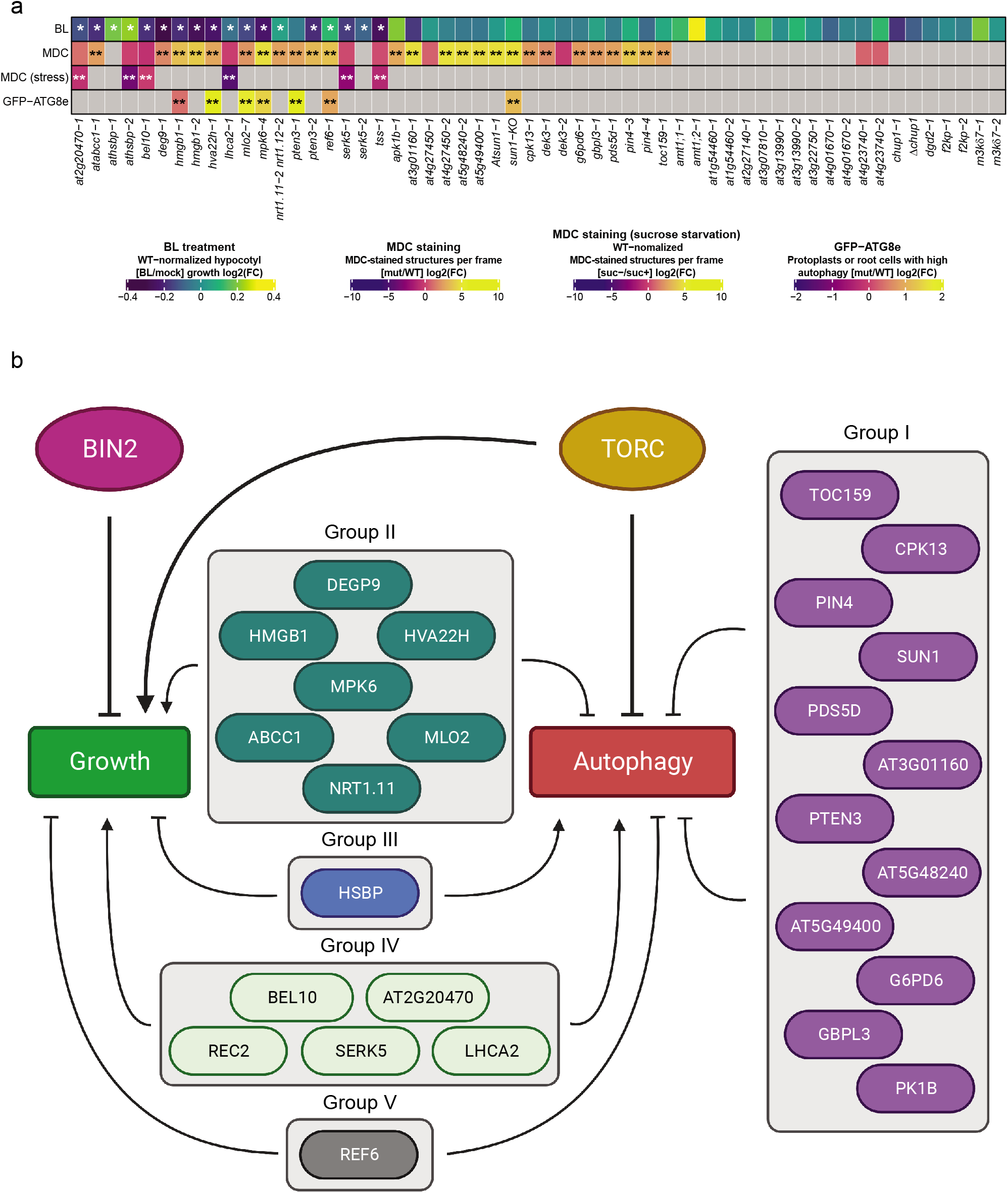
Identification of candidate genes with altered BR response and autophagy. **a**, (first row) Hypocotyl length response to BL treatment in selected mutant lines (four biological replicates each with n=6). Values shown are the log2 fold change in hypocotyl length (BL/Mock). * *p* <0.1, generalized linear model. (second and third row) Number of MDC-stained structures in selected mutants in normal growth conditions (second row, n=10) or sucrose starvation (third row, n=10). Values shown are the log2 fold change in the number of MDC-stained structures (mutant/WT, second row; -suc/+suc, third row). ** *p* < 0.05, two-sample t-test (fourth row) GFP-ATG8e expression in selected mutants. Values shown are the log2 fold change in the number of cells expressing GFP-ATG8e (mutant/WT). ** *p* < 0.05, two-sample t-test. **b**, Genes with significant response to BL or altered autophagy levels are organized into five groups according to their mutants’ phenotype.

In summary, we found a total of 26 genes out of the 42 selected candidates (61.9%) with a significant response to BR and/or altered level of autophagy (Supplementary Table 1). These results confirm the robustness of our pipeline as a way of selecting candidate proteins related to the brassinosteroid and/or autophagy pathways and, possibly, participating in plant growth/stress balance orchestrated by BIN2 and TORC.

## Discussion

Brassinosteroid and TORC have emerged as two key signaling pathways coordinating growth and stress responses. To gain a global view of BR and TORC crosstalk, we performed deep multi-omics profiling of the transcriptome, proteome, and phosphoproteome of *bin2* and *raptor1b* mutants. Our results provided evidence that BR and TORC pathways crosstalk at system-wide levels as they affect a common set of genes. Using the multi-omics data, we generated a kinase-signaling network, which was used to predict important regulators functioning in BR/TORC pathways. Our genetic studies indeed confirmed that many of the genes predicted from the network play important roles in BR-regulated growth and/or autophagy.

Our work supports previous transcriptome profiling studies of BR including (Wang et al., 2014; Kim et al., 2019; Liu et al., 2020) and TOR signaling (Ren et al., 2012; Xiong et al., 2013; Dong et al., 2015). Despite long-standing interest in BRs, comprehensive (phospho)proteomic profiles examining BR signaling are limited and proteome-wide identification of substrates of the key regulatory kinase BIN2 is lacking. By profiling *bin2* mutants, we identified transcripts, proteins, and phosphorylation sites whose proper expression is dependent on BIN2. Furthermore, using MAKS we provide a global catalogue of potential direct BIN2 substrates. In terms of TORC signaling, we substantially expand on the work of Salem et al., which provided an initial description of proteins that are mis-expressed in *raptor1b* (Salem et al., 2018) as well as the proteins and phosphorylation sites that respond to TOR inhibition via treatment with Torin 2, AZD8055, or rapamycin (Van Leene et al., 2019; Scarpin et al., 2020). Among our *raptor1b* 2,338 DE phosphosites, we found 46.8% (37/79) of the TOR-regulated phosphoproteins and seven of the TOR-interacting proteins from Van Leene et al, and 60.0% (48/80) of the Torin2 responsive phosphoproteins reported by Scarpin et al. Most importantly, through the generation and analysis of these multi-omics data we found a significant overlap of gene-products (i.e., transcript, protein, or phosphosites) that are regulated by both BIN2 and RAPTOR1B. Together our data suggest that a shared regulatory core exists between BR and TORC signaling.

Using our phosphoproteomics data, we inferred kinases that are differentially active in the analyzed mutant backgrounds. By assessing correlation between kinase activation and substrate phosphorylation levels, we reconstructed a kinase-signaling network predicting the regulatory linkages. Focusing on targets predicted by this regulatory network, and accounting for the phosphosites fold-change “directionality” on each mutant, we identified and tested a set of candidate genes for their involvement in BR/TORC signaling pathways. Mutants for forty-two genes were assayed and 26 of them showed an altered phenotype (Supplementary Table 1). To summarize these phenotyping results, we divided our gene set into 5 groups according to their different phenotypes in BR-regulated hypocotyl elongation and autophagy levels as a readout of TORC activity (Fig. 5b).

TOR signaling is known to be positively regulated by auxin and glucose availability (Xiong et al., 2013; Li et al., 2017; Schepetilnikov et al., 2017). Here, we found the auxin efflux carrier PIN4 and the chloroplast protein importer TOC159 as part of group I (autophagy negative regulators, Fig. 5b). This finding agrees with their corresponding functions: PIN4 creates an auxin sink at the cells below the quiescent center, a crucial event for normal auxin distribution (Friml et al., 2002). Additionally, TOC159 is essential for chloroplast biogenesis and, therefore, photosynthetic activity (Bauer et al., 2000). In parallel with this, we discovered homologs of proteins involved in autophagy and TOR signaling in human. Homo sapiens (Hsa) PTEN has been shown to negatively regulate both mTOR signaling and autophagy through independent pathways (Errafiy et al., 2013). Similarly, our results show Arabidopsis PTEN3 as an autophagy negative regulator (group I, Fig. 5). On the other hand, HsaHMGB1 can translocate to the cytoplasm and induce autophagy upon perception of reactive oxygen species (Tang et al., 2010). Here, HMGB1 is part of group II (positive BR regulator and negative autophagy regulation), suggesting an opposite function in plants.

Finally, our results identified previously unknown genes involved in BR and autophagy pathways. For example, mutant phenotype analysis indicated that MPK6 functions as a positive BR regulator and negative autophagy regulator. Although no direct role as a BR-induced growth has been established, MPK6 kinase is involved in a myriad of processes and has been shown to directly phosphorylate and activate BES1 to increase immune response (Kang et al., 2015). Furthermore, BIN2 can phosphorylate and inhibit MKK4, a direct MPK6 activator (Khan et al., 2013). Moreover, our signaling network prediction situates MPK6 as a possible RAPTOR1B regulator. Many other genes play a role in both BR-regulated growth and modulation of autophagy were identified by our analysis (Fig. 5b).

In summary, this study builds upon previous findings that BR and TORC crosstalk in the regulation of plant growth and stress responses (Zhang et al., 2014, 2016; Nolan et al., 2017b; Xiong et al., 2017; Salem et al., 2018; Vleesschauwer et al., 2018; Liao and Bassham, 2020; Nolan et al., 2020). Our multi-omics studies provide genome-wide evidence for extensive interactions between BR and TORC signaling pathways across three different regulatory layers (i.e., transcript, protein, and phosphorylation). We inferred a kinase signaling network that can effectively predict genes involved in these two pathways. Further functional studies should help refine the predicted regulatory network and define the mechanisms by which these regulators modulate both BR and TORC-regulated processes.

## Methods

### Plant material

*Arabidopsis thaliana* mutant lines *bin2-1* (Li and Nam, 2002), *bin2-3 bil1 bil2* (Yan et al., 2009), and *raptor1-1* (Anderson et al., 2005; Deprost et al., 2005) were used in this research as *bin2D, bin2T*, and *raptor1b*, respectively. The full list of seed stocks used in this work are summarized in Supplementary Data Set 12. All plants were grown in LC1 soil mix (Sungro) under long day conditions (16 h light, 22°C) unless said otherwise. Columbia-0 ecotype was used as wild-type control for all assays.

### QuantSeq library preparation and sequencing

Four biological replicates of 20-day-old rosette leaves were collected from WT and each mutant (*bin2T*, *bin2D*, and *raptor1b*) and immediately frozen in liquid N2. Tissue was ground for at least 15 minutes under liquid N2 using mortar and pestle. Total RNA was extracted using RNAeasy Plant Mini Kit with DNaseI treatment (Qiagen). Five-hundred ng of total RNA was used for QuantSeq 3’ (Moll et al., 2014) mRNA-seq library construction kit (Lexogen). Library sequencing was performed on an Illumina HiSeq3000. An average of 2,047,038 100-bp, single end reads was obtained per sample.

### Transcriptomic data analysis

Data processing followed the pipeline suggested by QuantSeq manufacturer. Briefly, reads were adapter- and quality-trimmed using BBDuk v37.36. Trimmed reads were mapped to the Arabidopsis reference transcriptome (TAIR10 annotation) using Star Aligner v2.5.3a (Dobin et al., 2013). Finally, transcript counts were extracted using HTSeq-count v0.11.2 (Anders et al., 2015). Differential expression was assessed using the PoissonSeq R package (Li et al., 2012). A q-value < 0.05 and absolute fold change > 1.32 was used as cutoff for designating differentially expressed transcripts. All data processing scripts were deposited in a github repository (see Data Availability section)

### Protein extraction for global proteome and phospho-proteome profiling

Three biological replicates from the same tissue collected for transcriptome analysis were used for protein extraction, using the urea-FASP method described by (Song et al., 2018a, 2018b, 2020) as follows: 250 mg of finely ground tissue was subjected to mechanical disruption using 250 mg ZrO_2_ beads in presence of 500 μL lysis buffer (8M urea; 100 mM TRIS-HCl, pH 7.0; 5mM TCEP) on a MiniG tissue homogenizer (Spex SamplePrep). Sample was clarified, and supernatant was transferred to a clean tube. Proteins were precipitated using rounds of 45-minute incubation at −80°C as follows: 1 round of ice-cold 100% acetone, 2 rounds of 80% acetone and 3 rounds of 0.2mM Na_3_VO_4_ in 100% methanol. Each round of incubation consisted of sample resuspension assisted by probe sonication, incubation at −20°C, centrifugation at 4,500xG for 10 minutes, and removal of supernatant. Extracted proteins were resuspended by sonication in urea resuspension buffer (URB, 8M urea in 50 mM TRIS-HCl, pH = 7.0; 5 mM TCEP; 1x phosphatase inhibitor cocktail) and further cleaned through Filter-Assisted Sample Preparation (FASP) using Amicon Ultra-4 30kDa MWCO filter units (Millipore) in the presence of UA buffer (8M urea in 100mM TRIS-HCl, pH = 8.0; 1x phosphatase inhibitor cocktail). Samples were reduced with 2mM TCEP, alkylated using 50 mM iodoacetamide (IAM) and digested into peptides using one round of overnight incubation at 37°C with 1:50 (enzyme:protein) trypsin (Roche, Cat. No. 03708969001) and a second round of incubation for 4 hours at 37°C with trypsin and Lys-C (Wako Chemicals, Catalog number 125-05061). Purified samples were further desalted using SepPack C18 columns (Waters). Tandem Mass Tag (TMT, Thermo Scientific) labeling was performed on 330 μg of purified peptides from each sample as previously reported (Song et al., 2020). TMT labeling reaction was stopped using 5% hydroxylamine and the quenched samples were then pooled. One hundred μg of labeled peptide was set aside for global proteome profiling and the remaining labeled sample was subjected to a second round of C18 desalting before phosphopeptide enrichment. Serial Metal Oxide Affinity Chromatography (SMOAC) method from Thermo Scientific was used for phosphopeptide enrichment. Briefly, High-Select TiO_2_ Phosphopeptide Enrichment kit (Thermo Scientific) was used as a first enrichment step and all flow-through was pooled, concentrated to almost dry on a SpeedVac and used as input for High Select Fe-NTA Phosphoptide Enrichment kit (Thermo Scientific). Phosphopeptides obtained from both enrichment processes were dried using a SpeedVac, resuspended in 0.1% formic acid in Optima grade H_2_O (Millipore) and pooled together.

### BIN2 Multiplexed Assay for Kinase Specificity

The MAKS as was performed based on the protocol described by (Jayaraman et al., 2017). Protein was extracted for MAKS from 1 g of leaf ground tissue from 20-days-old wild type Arabidopsis plants using the phenol-FASP protocol as described previously (Song et al., 2018b, 2020). Three mg of total purified protein was resuspended in URB, re-precipitated in ice-cold 100 mM NH_4_CH_3_CO_2_ in 100% methanol, and resuspended in kinase buffer (50 mM TRIS-HCl, pH = 7.7; 5 mM MgCl2; 5mM ATP; 1x phosphatase inhibitor cocktail). Resuspended protein was divided into 600 μg aliquots and incubated with either recombinant GST or GST-BIN2 at a 1:75 (enzyme:protein) ratio at 37°C with gentle shaking for 1 hour. After incubation, protein solution was subjected to FASP, reduced with 2mM TCEP, alkylated in 50mM IAM, and digested using trypsin as described by (Song et al., 2020). Three replicates were made for each treatment (i.e., GST and GST-BIN2). Two hundred μg of peptides from each replicate were used for TMT labeling. Phosphopeptide enrichment was performed on labeled peptides using SMOAC as described previously in this paper. Cloning of GST-BIN2 was described in (Yin et al., 2002); The fusion protein was purified using Glutathione agarose beads as described in (Jiang et al., 2019).

### LC-MS/MS

Chromatography was performed on an Agilent 1260 quaternary HPLC with constant flow rate of 600 nL min^−1^ achieved via a splitter. A Sutter P-2000 laser puller was used to generate sharp nanospray tips from 200 μm ID fused silica capillary. Columns were all in-house packed on a Next Advance pressure cell using 200 μm ID capillary. All samples were loaded into a 10 cm capillary column packed with 5 μM Zorbax SB-C18 (Agilent) and then connected to a 5 cm-long strong cationic exchange (SCX) column packed with 5μM PolySulfoethyl. The SCX column was then connected to a 20 cm long nanospray tip, packed with 2.5 μm C18 (Waters). For global protein abundance, 45 μg of labeled peptides and 24 ammonium acetate salt steps of 150 min each were used. For phosphoproteomics, 25 μg of enriched peptides and 15 salt steps were used. For MAKS, 30 μg of enriched peptides and 18 salt steps were used. Spectra were obtained using a Thermo Scientific Q-Exactive Plus high-resolution quadrupole-Orbitrap mass spectrometer on data-dependent mode as previously described (Zhang et al., 2019).

### Proteomics data analysis

Spectra for global protein abundance runs were searched using the Andromeda Search Engine (Cox et al., 2011) against the TAIR10 Arabidopsis proteome (https://www.arabidopsis.org/download_files/Proteins/TAIR10_protein_lists/TAIR10_pep_20101214) on MaxQuant software v1.6.1.0 (Tyanova et al., 2016) as reported previously (Zhang et al., 2019). Sample loading and internal reference scaling normalization methods were used to account for differences within and between LC-MS/MS runs, respectively (Plubell et al., 2017). Differential expression was assessed using the PoissonSeq R package (Li et al., 2012). A q-value < 0.1 was used as cutoff for designating differentially expressed proteins. Scripts for data analysis were deposited in a github repository (see Data Availability section).

### Phosphoproteomics data analysis

Spectra for both *bin2/raptor1b* mutant profling and MAKS were searched together using similar approach as with global protein abundance with exceptions. Briefly, MaxQuant software v1.6.10.43 was used instead and “Phospho (STY)” search for variable modifications was included. Sample loading and internal reference scaling normalization methods were used to account for differences within and between LC-MS/MS runs, respectively (Plubell et al., 2017). Differential expression was assessed using the edgeR R package (Robinson et al., 2010). A q-value < 0.1 was used as cutoff for designating differential phosphorylation. See Data Availability section for the full analysis script.

### Motif enrichment analysis

Motif enrichment was performed using the motifeR R package (Wang et al., 2019) with default settings, serine or threonine as the central residues, a p-value threshold of 0.001, and TAIR10 protein annotation as background reference. Enrichment p-value was calculated using hypergeometric testing.

### Analysis of overlap between BIN2 MAKS and *bin2* mutant datasets

To find overlapping phosphosites, we defined any two distinct phosphosites as identical if they originated from the same phosphoprotein and were less than 10 amino acid residues apart. This approach was used to account for cases where phosphosites were not localized to a specific amino acid on a given peptide. Overlap statistical significance was assessed by hypergeometric test.

### Kinase activation loop prediction

Protein kinases were identified using a modified version of the pipeline described by (Walley et al., 2013). Briefly, all 35,386 protein sequences available in the TAIR10 annotation (https://www.arabidopsis.org/download_files/Proteins/TAIR10_protein_lists/TAIR10_pep_20101214) were searched for kinase domain using The National Center for Biotechnology Information batch conserved domain search tool (Lu et al., 2020). From this list of 1,522 proteins with identified kinase domain, 878 were also annotated with activation loop (A-loop) coordinates by the search tool. The kinase domains of proteins lacking the A-loop coordinates were aligned using MAFFT (Katoh and Standley, 2013) and the well conserved A-loop beginning (DFG) and end (APE) motifs were manually searched. An extra 482 A-loop coordinates were obtained, for a total of 1,360 protein kinases with A-loop coordinates.

### Kinase-substrate network

Kinases with differential phosphorylation inside the A-loop (activated kinases) were used as regulators to build the kinase-substrate network. For this, the Spearman and Pearson correlation between a regulator and the rest of differentially phosphorylated peptides was calculated as described previously (Walley et al., 2013).

### BL response assays

Seeds were vapor-phase sterilized in chlorine gas, stratified at 4°C for 1 week, and germinated in half-strength Linsmaier and Skoog media (Caisson Labs, catalog number LSP03) supplemented with either DMSO or 100 nM brassinolide (BL). Seedlings were grown for 7 days at 22/18°C (day/night), 16 hours of light, 40% relative humidity, and light intensity of 120 μmol m^−2^ s^−1^. Seedlings were imaged and hypocotyl length was measured using Fiji software (Schindelin et al., 2012). A generalized linear model with treatment and genotype as factors and controlling for random effects of replicate and plate was applied and a threshold of “genotype by treatment interaction” *p*-value < 0.1 was set as significance cutoff. Four replicates were made with six seedlings being measured for each replicate. The whole experiment was repeated at least two times per genotype.

### MDC staining

Seven-day-old Arabidopsis seedling roots were stained with monodansylcadaverine (MDC, Sigma) as described previously (Contento et al., 2005). Cells within the root elongation zone for ten seedlings were observed and imaged using a Zeiss Axio Imager.A2 upright microscope (Zeiss) equipped with Zeiss Axiocam BW/color digital cameras and a DAPI-specific filter at the Iowa State University Microscopy and Nanoimaging Facility. The number of MDC-labeled puncta in each image was counted and averaged from at least 10 images per sample. For sucrose starvation, 7-day-old seedlings grown on solid ½ MS medium were transferred to solid ½ MS medium lacking sucrose and kept in darkness for an additional 3 days before staining. Significance was assessed through two-sample t-test for MDC staining under normal growing conditions, whereas a generalized linear model with treatment and genotype as factors and controlling for random effects of replicates was used for MDC staining under sucrose starvation. A p-value < 0.05 was used a cutoff on both cases.

### GFP-ATG8e protoplast assay and stable lines

Protoplasts were extracted from leaves from 20-day-old plants and transformed as described previously (Wu et al., 2009). Protoplasts were observed on a LSM-700 confocal microscope (Zeiss) using a FITC filter, and protoplasts with more than three visible autophagosomes were counted as active for autophagy as previously described (Yang et al., 2016; Pu et al., 2017). One hundred protoplasts were analyzed per genotype and the experiment was repeated three times. Homozygous mutant lines were crossed to plants expressing GFP-ATG8e and stable lines were obtained. Roots from GFP-ATG8e-expressing lines that were also homozygous for the mutation were analyzed on LSM-700 confocal microscope (Zeiss) using a FITC filter. Cells within the root elongation zone of no less than fifteen seedlings were photographed and the number of autophagosomes in each image was counted and averaged from at least 10 images per sample. Significance was assessed in both cases by two-sample t-test and a *p*-value < 0.05 was used as cutoff.

### Data Availability

The RNA-seq data generated by this work has been deposited at the National Center for Biotechnology Information (NCBI) Short Read Archive (SRA) as BioProject accession number PRJNA678744. The original MS proteomics raw data, as well as the MaxQuant output files, can be downloaded from MassIVE (http://massive.ucsd.edu) using the identifier MSV000086460 for *bin2D, bin2T*, and *raptor1b* mutants profiling and MSV000086462 for the BIN2 MAKS. All the scripts used for this work are available on the following github repository: https://github.com/chrisfmontes/BIN2_RAPTOR1B_MULTIOMICS.

## ACKNOWLEDGEMENTS

This work was supported by the Iowa State University Plant Science Institute (YY and JWW), NIH R01GM120316 (YY, DB, JWW), NSF IOS-1818160 (YY and JWW), and USDA NIFA Hatch project IOW3808 funds to JWW. NMC is supported by a USDA NIFA Postdoctoral Research Fellowship (2019-67012-29712) and TMN is supported by the National Science Foundation Postdoctoral Research Fellowships in Biology Program (Grant No. IOS-2010686).

This manuscript was formatted in Overleaf using the Henriques Lab bioRxiv template.

---

**Supplementary Fig. 1.**
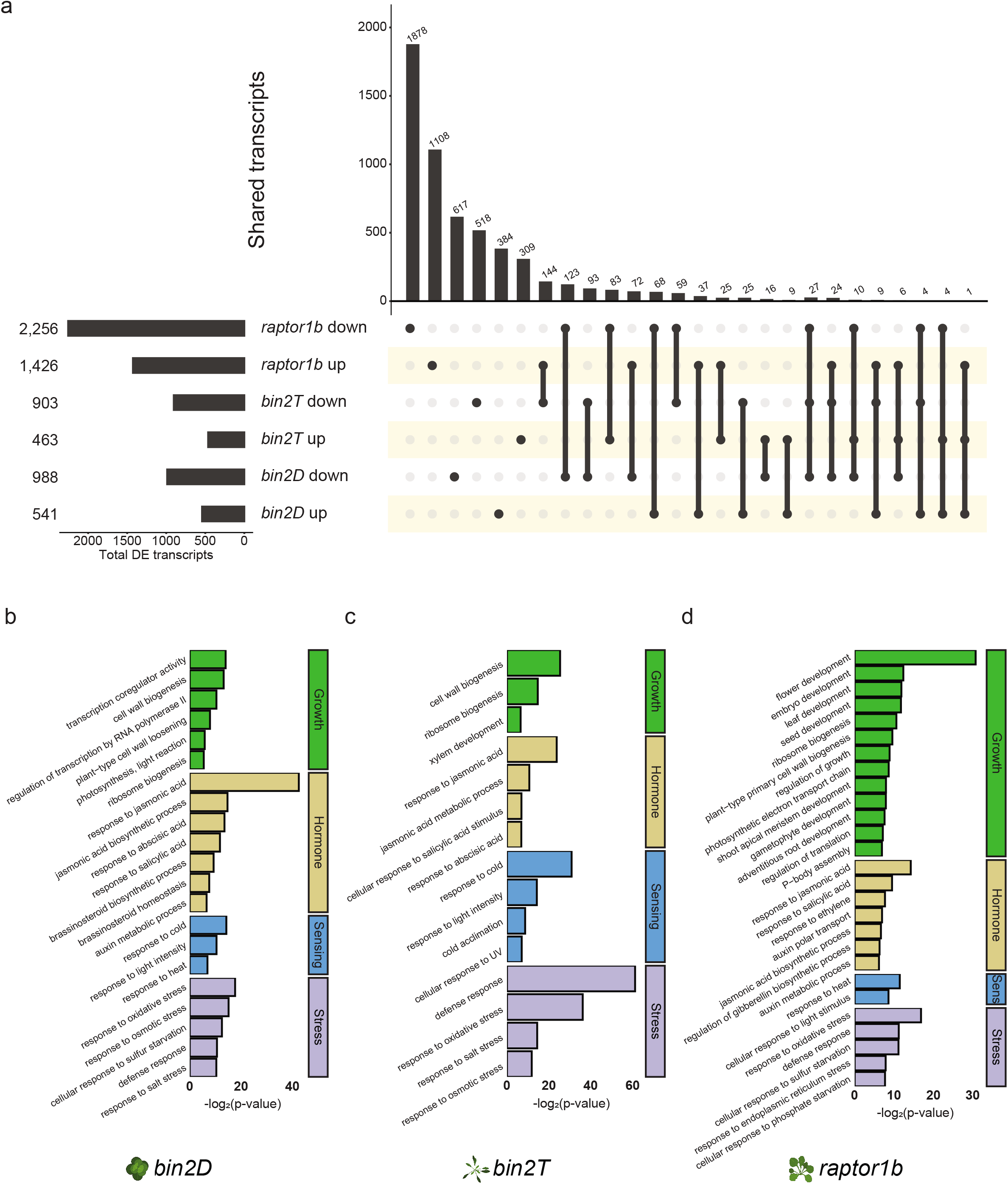
Transcriptomic analysis of *bin2* and *raptor1b* mutants. **a**, UpSet plot showing overlap of DE transcripts between *bin2D, bin2T*, and *raptor1b*. **b-d**, Selection of significant GO biological processes among differential expressed transcripts on *bin2D* (b), *bin2T* (c), and *raptor1b* (d). Differential expression was defined as q-value <0.05 and absolute fold-change > 1.32.

**Supplementary figure 2.**
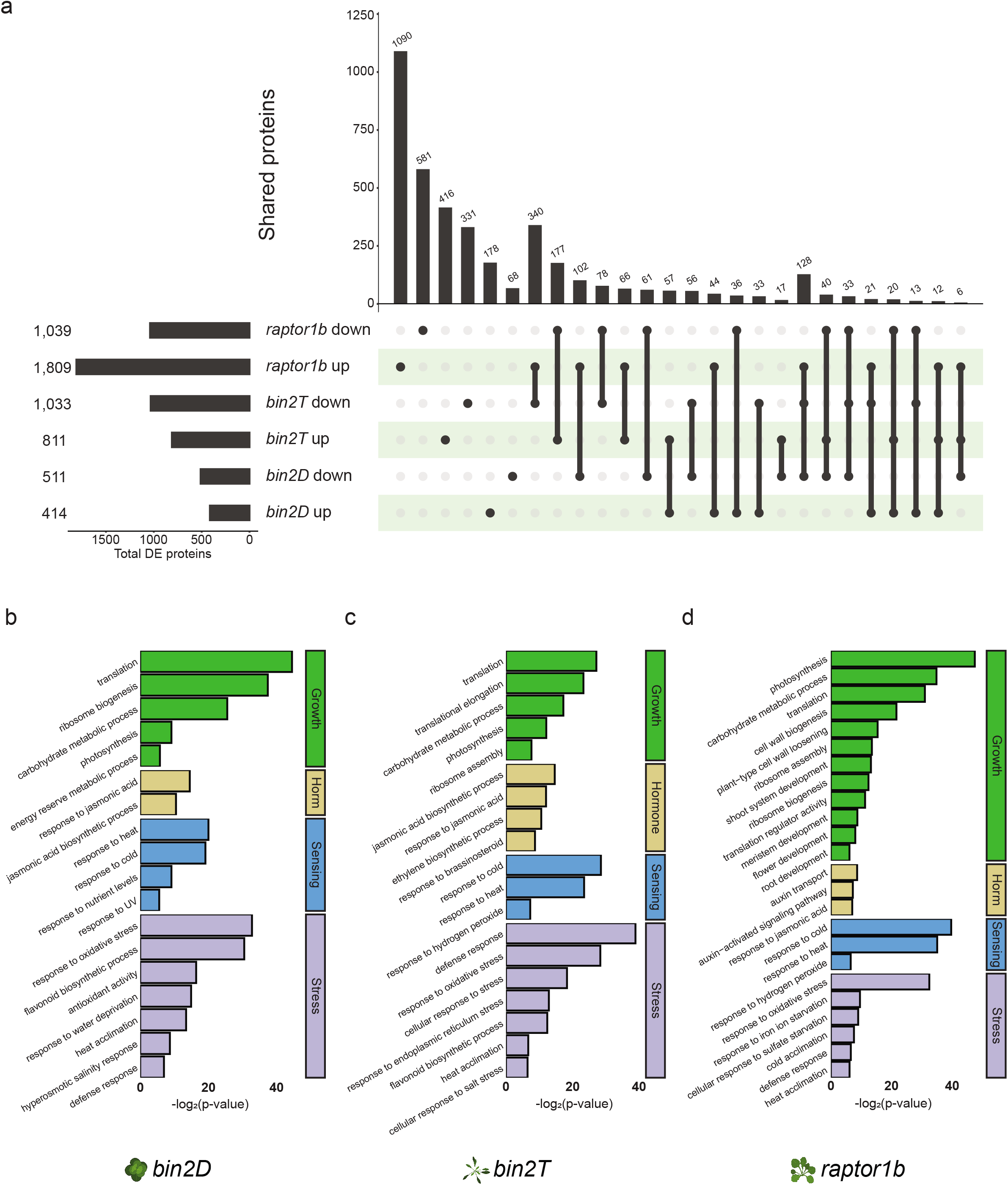
Proteomic analysis of *bin2* and *raptor1b* mutants. **a**, UpSet plot showing overlap of DE proteins between *bin2D, bin2T*, and *raptor1b*. **b-d**, Selection of significant GO biological processes among differential expressed proteins on *bin2D* (b), *bin2T* (c), and *raptor1b* (d). Differential expression was defined as q-value <0.1.

**Supplementary figure 3.**
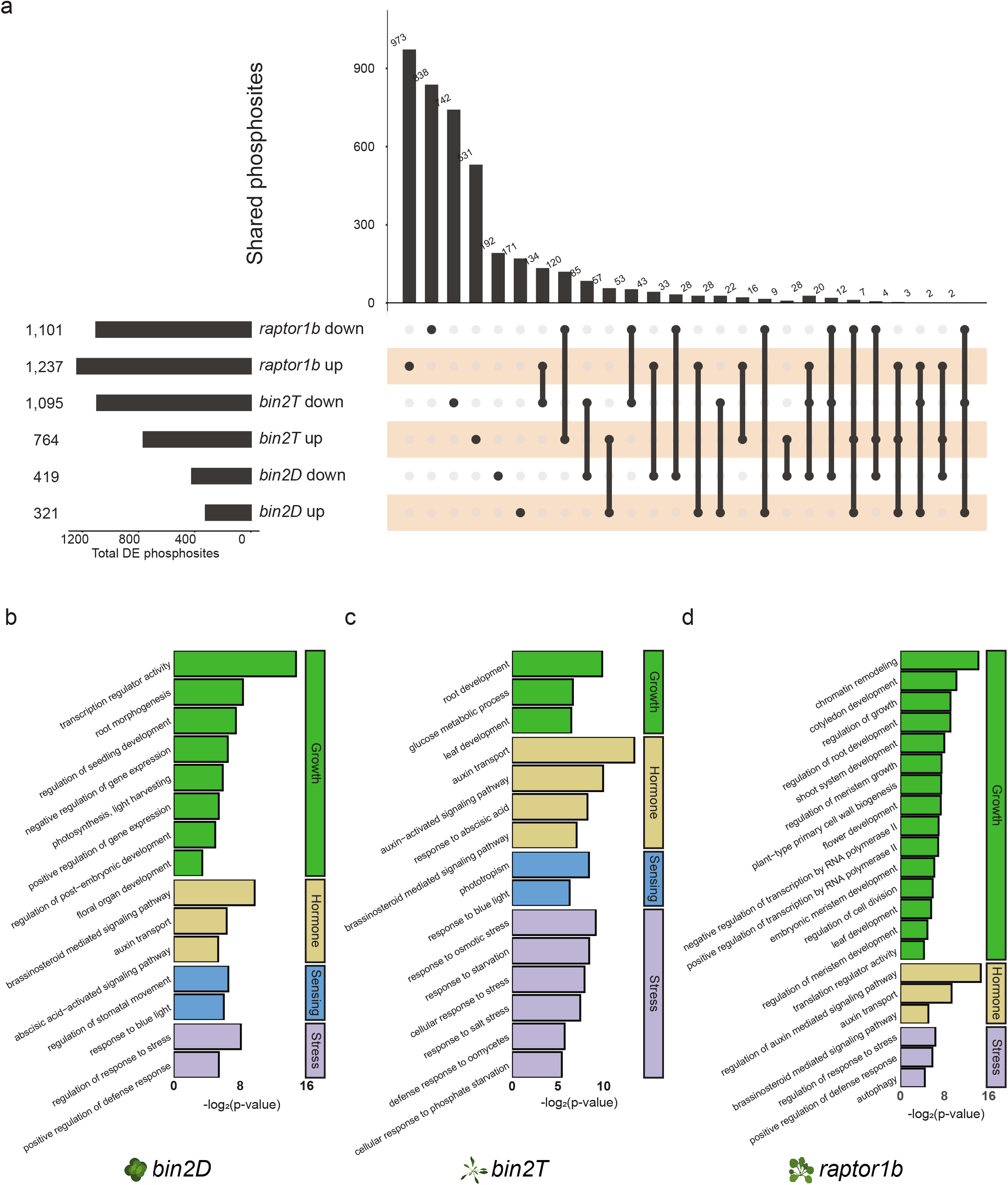
Phosphoproteomic analysis of *bin2* and *raptor1b* mutants. **a**, UpSet plot showing overlap of differentially regulated phosphosites between *bin2D, bin2T*, and *raptor1b*. **b-d**, Selection of significant GO biological processes among differentially upregulated phosphosites on *bin2D* (b), differentially downregulated phosphosites on *bin2T* (c), and differentially downregulated phosphosites on *raptor1b* (d). Differential phosphorylation was defined as q-value <0.1.

**Supplementary figure 4.**
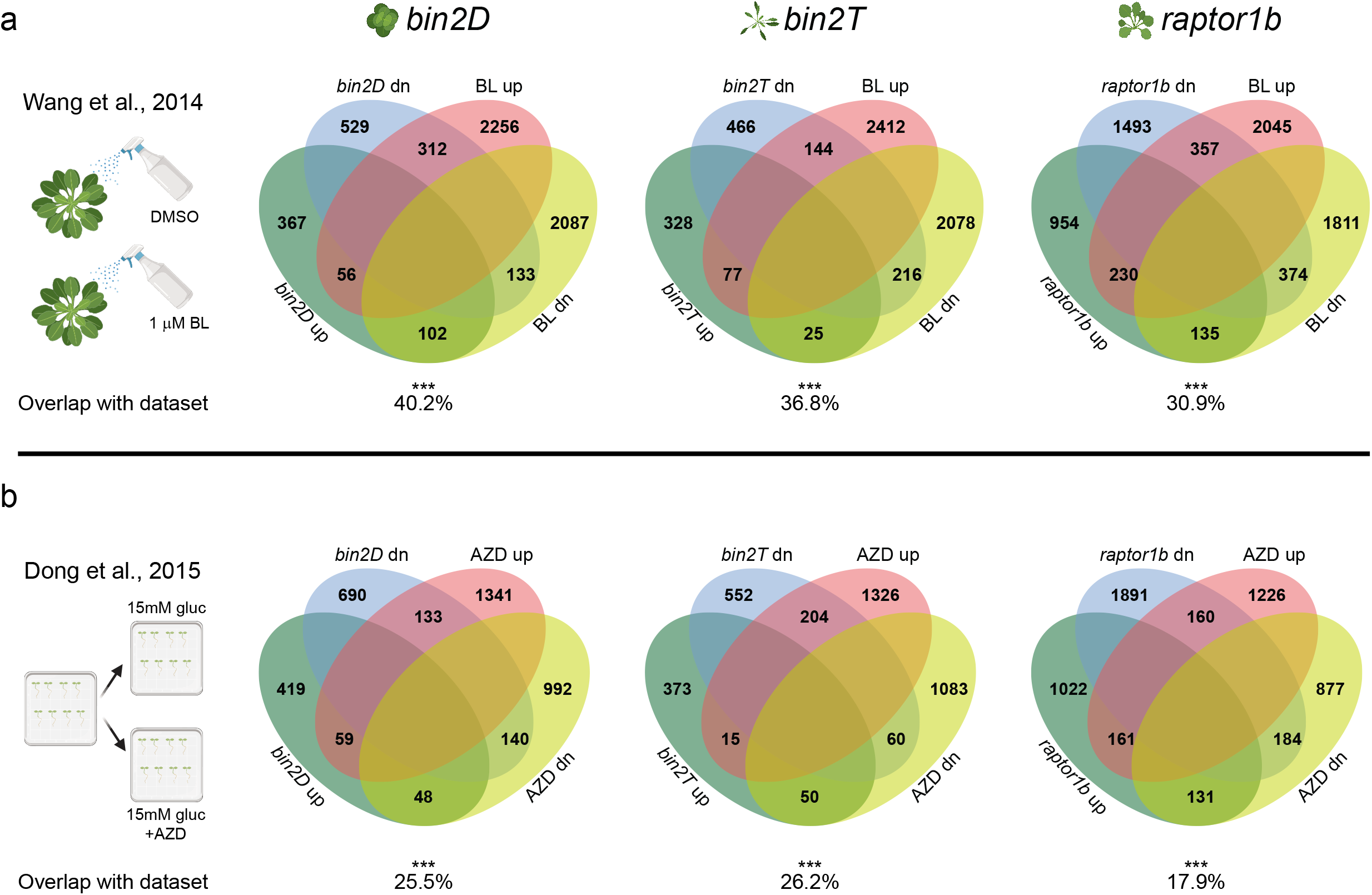
Overlap with previously reported datasets. **a**, **b**, Overlap between DE transcripts in *bin2D, bin2T*, and *raptor1b* and BL-induced DE transcripts from Wang *et al*. (a) or AZD-induced DE transcripts from Dong *et al*. (b). Overlapping significance was assessed using hypergeometric test. *** *p* < 0.001

**Supplementary figure 5.**
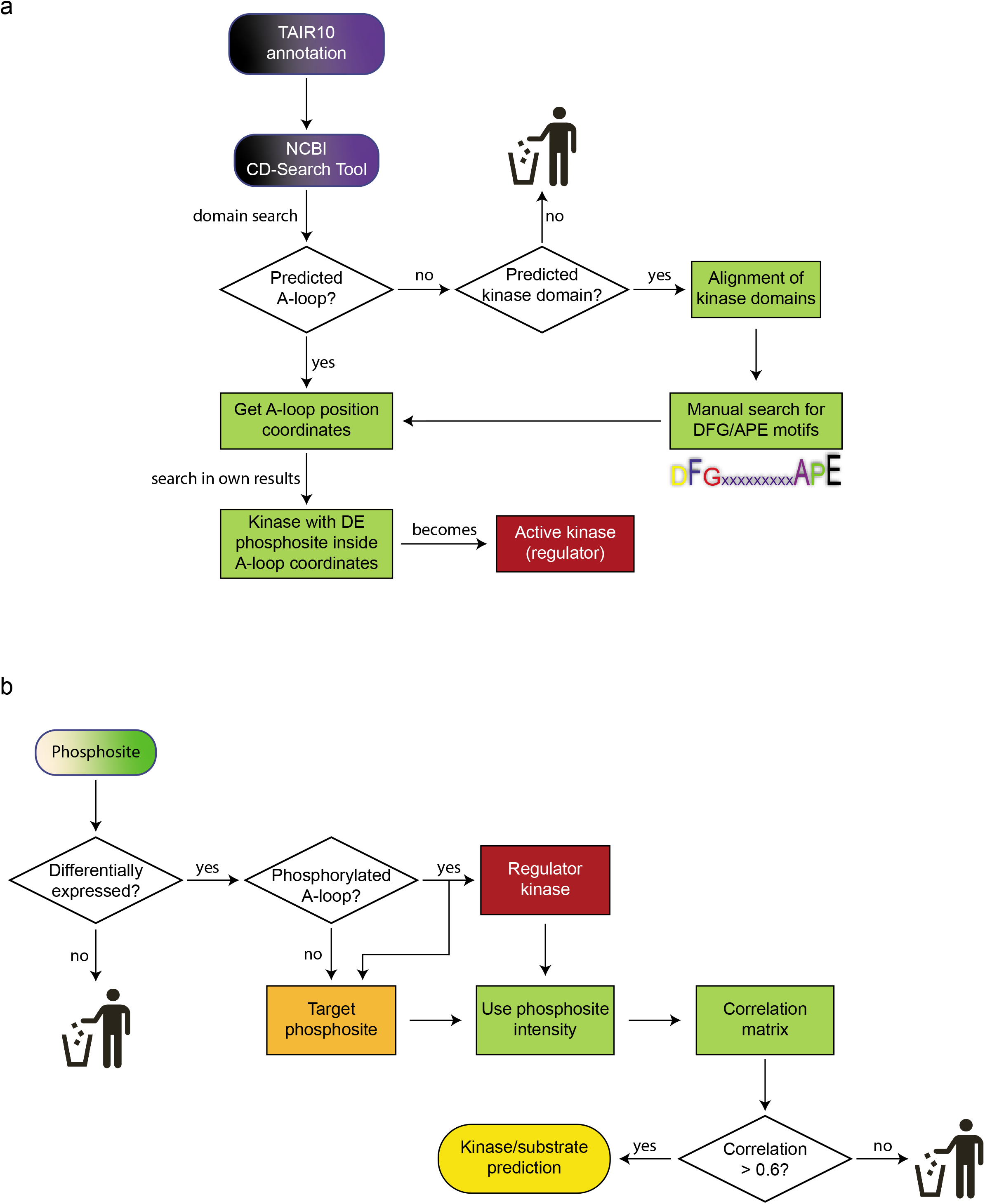
Workflow of signaling network reconstruction. **a**, Pipeline for activation loop coordinates annotation for Arabidopsis kinases. **b**, Logic diagram for kinase/substrate prediction.

**Supplementary figure 6.**
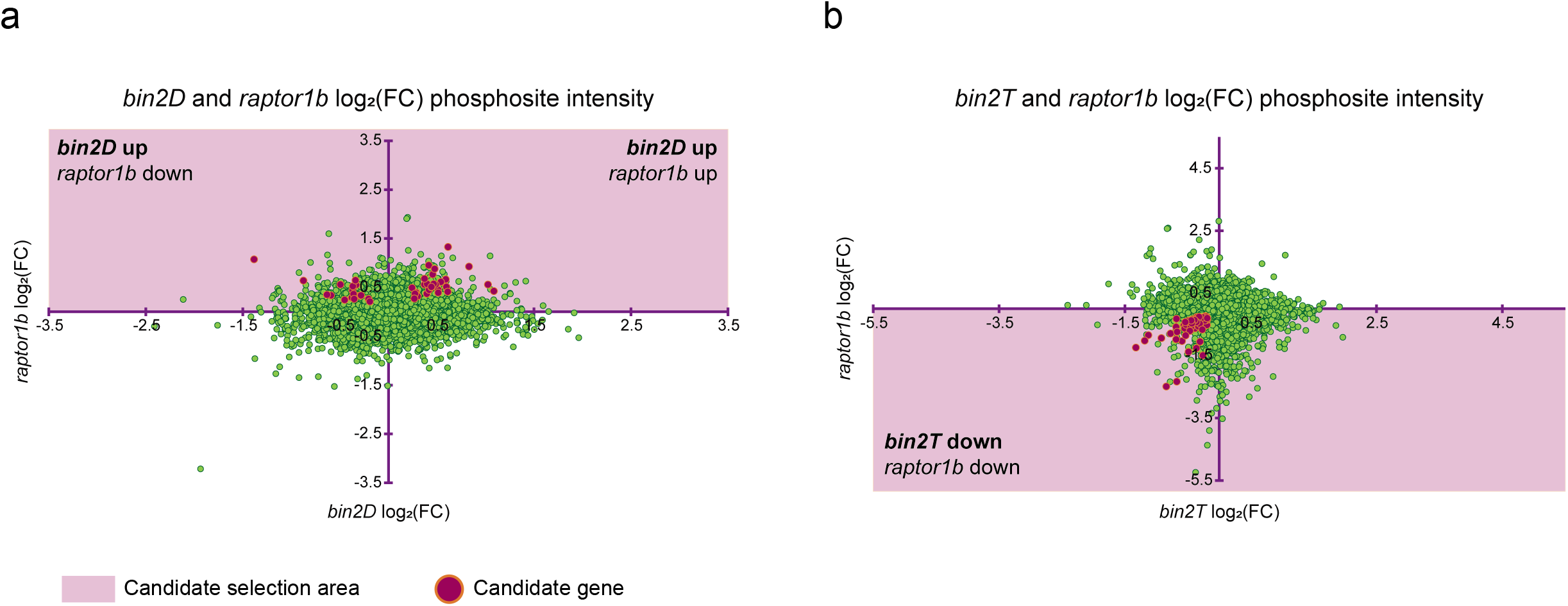
Phosphosite intensity fold-change scatterplots and selection of candidate genes. **a**, **b**, Scatterplot showing phosphosite intensity log_2_ fold-change (mutant/WT) for *raptor1b* on the x-axis and *bin2D* (a) or *bin2T* (b) on the y-axis. Red dots indicate phosphosites from selected candidate genes.

**Supplementary table 1.** BIN2/RAPTOR1B responsive candidate genes screening results

|  | No. different genes | % |
| --- | --- | --- |
| Candidate genes assayed by mutant screening | 42 | 100 |
| Predicted as target by network | 42 | 100 |
| Predicted as activated kinase | 3 | 7.1 |
| Predicted as BIN2 target by network | 17 | 40.5 |
| Total genes with mutant(s) showing consistent BR or autophagy altered response | 26 | 64.2 |
| Genes with mutant showing altered BR response | 14 | 33.4 |
| Genes with mutant showing altered autophagy response | 26 | 64.2 |

## Notes

### Competing Interest Statement

The authors have declared no competing interest.

